# Empathy Accelerates Subjective Time Perception, Independent of Physiology but Not of Visual Attention

**DOI:** 10.64898/2025.12.25.696484

**Authors:** Soroosh Golbabaei, Mina Hosseinnezhad, Zahra Karami, Khatereh Borhani

**Affiliations:** Department of Psychiatry and Psychotherapy, Jena University Hospital, Jena, Germany; German Center for Mental Health (DZPG), partner site Halle-Jena-Magdeburg, Jena, Germany; Institute for Cognitive and Brain Sciences, Shahid Beheshti University, Tehran, Iran; Laboratory for the Experimental Research of Religion, Masaryk University, Brno, Czech Republic

**Keywords:** Empathy, Time Perception, Physiological Signals, Eye tracking, Alexithymia, Autism Spectrum Disorder

## Abstract

Recent work suggests that empathic engagement can influence cognitive and perceptual processes, yet its impact on time perception remains unclear. Whereas internal clock models would predict that empathy-induced physiological arousal lengthen perceived duration, other frameworks offer opposite predictions. Specifically, findings that empathy reduces perceived distance, combined with construal level, and motivational dimensional theory, together predict that empathy accelerate the subjective passage of time. Moreover, although eye-movements are crucial in empathy, it remains unclear how visual attention modulates time perception during empathic experiences. Thus, we employed an empathy-for-pain and duration estimation task, while measuring participants’ heart rate, skin conductance, and eye-movements, to investigate how empathy and physiological arousal influence time perception. Our findings show that empathy leads to a faster perceived passage of time. Despite increases in physiological arousal during empathy, arousal was not associated with changes in time perception. Instead, greater visual attention to the eyes predicted stronger temporal contraction. Furthermore, trait empathy and autism quotient were related to this effect, whereas alexithymia was not. These results provide the first empirical evidence that empathy alters time perception, elucidate the contribution of physiological and visual mechanisms to this phenomenon and extend current theories of time perception and empathy.

---

Empathy has long captivated scientific interest due to its fundamental role in social communication. Over the past decades, researchers have sought to define empathy more precisely and examine its relationship with a broad range of social and psychological phenomena (Cuff et al., 2014; Gibbons, 2011; Hall & Schwartz, 2019; Zaki, 2017). Empathy is widely conceptualized as a multifaceted construct encompassing both affective empathy, the capacity to share others’ emotional states, and cognitive empathy, the ability to understand others’ emotions and perspectives (Cuff et al., 2014; Reniers et al., 2011; Walter, 2012).

These components have been linked to diverse social and behavioral outcomes, such as prosocial behavior (Li et al., 2024; Morelli et al., 2014), aggression (Björkqvist et al., 2000; Li et al., 2024), and morality (Decety & Cowell, 2014; Samani et al., 2022) and clinical and subclinical conditions including autism spectrum disorder (Bird & Viding, 2014; Golbabaei, et al., 2022; A. Smith, 2017) and alexithymia (Bird & Viding, 2014; Golbabaei, et al., 2022; Grynberg et al., 2010). Yet, just recently, as research has turned toward embodied aspects of empathy (Walsh, 2013; Wilson, 2002) and shared representation (Lamm et al., 2016; Singer et al., 2004) a growing body of research has begun to focus on the role of self–other distinction in empathic processes. This line of work suggests that empathy extends beyond social behavior, rather it also involves changes in bodily states and signals (Bukowski et al., 2020; Golbabaei, et al., 2022; Hasan, et al., 2025; Hasan, et al., 2025; Lamm et al., 2016), self-perception (Lamm et al., 2016; Lopez et al., 2013), and even self-other boundaries(Perry et al., 2015; Yang et al., 2024). Such changes may in turn influence a variety of cognitive and perceptual domains, including how we perceive space (Golbabaei & Borhani, 2024), our own body (Martínez-Pernía et al., 2023; Weiler et al., 2024), and potentially, time.

Humans generally possess a reliable sense of time across. Yet, subjective time perception is remarkably flexible, influenced by attention (Yan et al., 2025), arousal (Smith et al., 2011a), emotional valence (Droit-Volet et al., 2004), salience (Wittmann, 2015), rhythm of environmental events (Droit-Volet & Wearden, 2002) and more importantly by how we perceive the world around us and ourselves within the environment (Wittmann, 2015). It is therefore plausible that empathy, by altering our sense of self and our relation to others, could also modulate the perception of time. Closely related to empathy, effect of emotions on temporal judgments have been recently studied widely. Early studies suggested that emotionally charged stimuli, especially aversive (Dirnberger et al., 2012) or high-arousal ones (Drew et al., 2003) lead to temporal dilation, such that time seems to pass more slowly. For example, participants perceived happy or angry faces as lasting longer than neutral faces when imitation was allowed (Droit-Volet et al., 2011). Likewise, inducing sadness led to an expansion of subjective time (Benau & Atchley, 2020). These findings were traditionally interpreted through the lens of the internal clock model, positing that heightened arousal accelerates a pacemaker-like mechanism, increasing temporal pulses. This increase is reflected in elevated physiological activity such as higher heart rate, and ultimately leads to lengthening of perceived time (Gibbon, 1977; Gibbon et al., 1984).

Based on earlier findings, one might predict that empathy would similarly dilate perceived time. Because empathic engagement involves sharing another’s emotional state, it seems reasonable to assume that it would produce effects comparable to those observed when directly experiencing emotions, namely a lengthening of subjective duration. However, later research has revealed that the relationship between emotion and time perception is far more complex, varying across emotional categories and experimental contexts (S. D. Smith et al., 2011b). Not all emotions lead to time dilation, and in some cases, emotionally charged states can even shorten perceived duration (Colonnello et al., 2016; Gil & Droit-Volet, 2011; Moreira et al., 2025; Weng et al., 2022). For example, Kliegl et al., (2015) reported that sad and neutral images did not differ in their influence on subjective time perception. Interestingly, a recent study found that participants perceived both pleasant and unpleasant images as lasting shorter than neutral images (Moreira et al., 2025).This inconsistency suggests that factors beyond arousal, such as motivations, changes in self-representation and embodiment, may contribute to temporal distortions.

A more recent theory of time perception, the motivational dimensional model of time perception (Gable et al., 2016, 2022; Gable & Poole, 2012), has criticized the previous approach of arousal-valence in time perception and has proposed that motivation is a key factor in time perception, where approach motivation accelerates time perception and avoidance motivation does the opposite. For example, when participants experienced an approach motivation, irrespective of emotions and arousal, time flied. Moreover, when approach motivation was experimentally manipulated, time passed faster (Gable & Poole, 2012). Besides, approach-oriented emotions, namely sadness and anger have been associated with shorter perceived durations, whereas disgust, which is motivationally avoidance-oriented has led the time to pass slower. In addition, when motivation has been experimentally manipulated, higher approach motivated sadness and anger has led to faster passage of time(Gable et al., 2016).

Empathy extends beyond mere emotion sharing and is inherently dyadic (Zaki et al., 2008), rooted in social and evolutionary functions that promote prosocial motivation, helping behavior, and the alleviation of others’ distress (Molnar-Szakacs, 2011; Pavey et al., 2012; Samani et al., 2022; Zaki, 2014). Neuroimaging research shows that empathy recruits somatosensory cortical regions involved in representing bodily states and elicits subtle muscle activations, consistent with embodied resonance mechanisms that prepare individuals for action (Molnar-Szakacs, 2011). Thus, empathy is related to motivation to approach the target to provide help, and alleviation of pain. Therefore, based on the motivational dimensional model (Gable et al., 2016, 2022; Gable & Poole, 2012), empathy should accelerate the passage of time.

More importantly, empathy involves reduced self–other distinction and a partial overlap of bodily and neural representations between observer and target. This self–other overlap, by altering perception of the self in relation to the environment, influences fundamental perceptual processes (Chambon et al., n.d.; Farmer & Maister, 2017; Golbabaei & Borhani, 2024). A recent work demonstrated that empathy reduces perceived physical distance between observer and target, suggesting that empathic experience compresses perceptual space (Golbabaei & Borhani, 2024). According to construal level theory, psychological distance is organized along multiple dimensions, including spatial, temporal, social, and hypothetical, that are interrelated and egocentrically anchored to the self. Decreasing distance along one dimension can lead to a reduction along others, including temporal distance (Fiedler et al., 2012; Kanten, 2011; Rim et al., 2009; Trope & Liberman, 2010). Consequently, if empathy reduces perceived spatial (Golbabaei & Borhani, 2024) and social distance (Bilali et al., 2018; Méndez Fernández et al., 2022) by blurring self–other boundaries, it may also compress perceived time. Such an effect would contrast with the predictions of the internal clock model, emphasizing instead how changes in embodiment and self–other representation can shape temporal experience.

Visual attention also plays a key role in both empathy and time perception. Empathic responses are associated with increased attention to emotionally salient facial features, particularly the eyes, and reduced attention to less socially informative regions such as the mouth (Cowan et al., 2014; Golbabaei, et al., 2023). If empathy alters time perception, these shifts in visual attention may also contribute to temporal distortions in empathic contexts. However, the contribution of differential gaze patterns toward facial features during empathic engagement remains unexplored.

Finally, interindividual differences in empathy-related personality traits may further influence temporal experience. Trait empathy, the stable tendency to empathize with others, differs from context-dependent state empathy, though the two are closely related. Moreover, traits such as alexithymia and autistic characteristics are negatively associated with empathic capacity (Golbabaei, et al., 2022, 2023; Grynberg et al., 2010; Shamay-Tsoory et al., 2002; Smith, 2017). Investigating how these traits modulate time perception during empathic engagement could provide insight into the interplay between empathy, embodiment, and temporal cognition.

To address these questions, we employed an empathy-for-pain paradigm in which participants reported perceived passage of time across conditions eliciting varying levels of empathic engagement, including high empathy, low empathy and non-empathic attentional demanding condition. Heart rate and electrodermal activity were recorded to index physiological arousal. If time perception were determined primarily by emotional arousal, as predicted by the internal clock model, higher arousal should lengthen perceived duration. In contrast, our main hypothesis was that empathy alters time perception through shared representation, reduced self–other distinction, and approach motivations, leading to temporal compression, independent of or inversely related to physiological arousal. Moreover, we hypothesized that greater visual attention to the eyes and less visual attention to lips would be associated with stronger temporal compression, and that individuals with higher trait empathy and lower alexithymia and autistic traits would exhibit stronger modulation of time perception during empathic engagement.

## Method

### Participants

A power analysis (Faul et al., 2007) indicated that 42 participants were required for a paired *t*-test (95% power, *d*=0.5, *α*=.05). To allow for correlational analyses and potential data loss in physiological and eye-tracking measures, 70 participants were recruited (42 females; *Mean*_age_=25.72, *SD_age_*=3.07) via social media and in-person invitations. All participants reported no neurological or physiological disorders, were medication-free, and had normal or corrected-to-normal vision. Written informed consent was obtained, and the study was approved by the Ethics Committee of Shahid Beheshti University.

### Procedure

Three days prior to the experiment, participants completed an online survey including the IRI, TAS-20, AQ, and demographic, health, and medication questions. On the day of testing, they were invited to the laboratory and seated comfortably in front of a laptop equipped with an eye tracker, where instructions were delivered verbally. ECG and EDR electrodes were then attached, and after signal stabilization, participants positioned their chin on a chin rest with viewing distance adjusted to approximately 65 cm. Eye-tracker calibration followed, after which participants performed the empathy-for-pain and duration estimation task (see *Empathy-for-Pain and Duration Estimation Task*).

### Stimuli

Four validated empathy-evoking images (two male, two female) depicting facial expressions of pain induced by electrical stimulation to the hand were used. These stimuli were adopted from a previously published dataset and have been shown to reliably elicit empathy (Mosalmannejad et al., 2025; Zarei et al., 2023). Each image was overlaid with an oval mask (Figure 1A).

**Figure 1.**
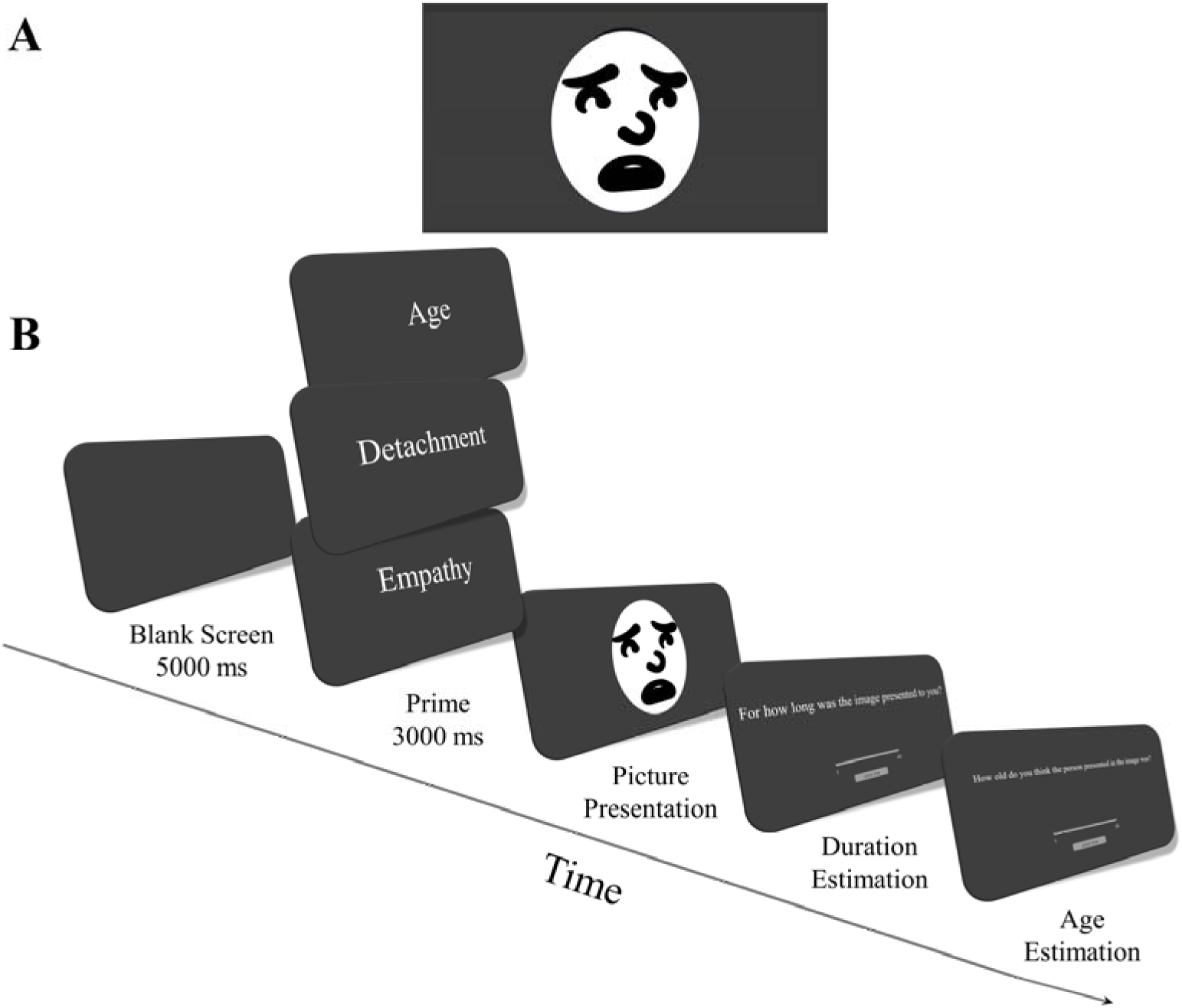
Schematic Diagram of the Empathy for Pain and Duration Estimation Task. Notes. A: a sample of pictures used in the study B: schematic diagram of the task. Pictures were presented for 10, 12, or 16 seconds. There was no time limit for responding to duration estimation and age estimation questions. Facial expressions were replaced by cartoon illustrations in accordance with repository restrictions.

### Empathy-for-Pain and Duration Estimation Task

Each trial began with a 5-s blank screen, followed by a 3-s cue word (empathy, detachment, or age) indicating the trial condition (high empathy, low empathy, or age estimation). The cue instructed participants to empathize with the person and put themselves in their place, remain indifferent and emotionally detached, or estimate the person’s age during subsequent image viewing. The use of the age condition serves as an additional attentional demanding control and it is hypothesized that the results for this condition will align with and be a stronger version of the detachment condition. An image depicting a person in pain was then presented. Images were shown for 10, 12, or 16 s under each condition (9 presentations per image). After viewing each image, the participant answered four questions without time limit. They first estimated the image duration using a visual analog scale (1–40 s), followed by age estimation on a visual analog scale (1–35 years). The Age estimation question served to mask the study’s true objective and to equate attentional demands across conditions. In total, the task comprised 36 trials (4 images × 3 durations × 3 conditions). A schematic diagram of the task is shown in Figure-1B.

To assess cognitive and affective empathy, participants completed a post-task rating procedure following established protocols (Golbabaei & Borhani, 2024; Pfabigan et al., 2015). Each picture was presented twice, once in the high-empathy condition and once in the low-empathy condition, for 5-s per presentation. After each presentation, participants rated their cognitive and affective empathic responses using a slider without time constraints.

Cognitive empathy was assessed with the question, “How unpleasant was the stimulus for the other person?”, whereas affective empathy was measured with the question, “How unpleasant was it for you to see the other person receive the stimulus?”. Responses were recorded on a 7-point Likert scale ranging from 1 (not at all) to 7 (extremely). The task was coded using PsychoPy.

### Physiological Measurement and Analysis

Electrocardiogram (ECG) data were recorded at 250 Hz using a Bioline acquisition system, with electrodes placed on the wrist and clavicle. ECG signals were band-pass filtered between 1 and 50 Hz using the PhysioData-Toolbox. R-peaks were initially detected automatically and subsequently reviewed by two independent experts to identify missed or incorrectly detected peaks, which were corrected manually. Image presentation periods for each condition were defined as epochs, and heart rate and interbeat intervals were extracted by averaging values across trials within each condition.

Skin conductance activity was recorded using the same device, with electrodes placed on the index and middle fingers of the non-dominant hand. The signal was low-pass filtered at 2 Hz and detrended prior to peak detection. Skin conductance responses (SCRs) were identified using a valley-to-peak method with a minimum amplitude of 0.02 µS and a minimum rise time of 0.5 s. To account for overlapping SCRs, inflection points were detected by computing and smoothing the first derivative of the signal with a 0.5 s window. Negative-to-positive zero crossings were treated as inflection point, marking the peak of the preceding SCR and the onset of the subsequent one. The total number of SCRs within each epoch was calculated and used. Physiological signals could not be obtained for nine participants. ECG data from five participants were excluded because of excessive noise, and EDR data from three participants were excluded due to a lack of physiological responsiveness.

### Eye-tracking Analysis

We used an SMI-Red-250 to record eye-movements. Calibration employed a 13-point procedure, followed by 5-point validation. Eye-movement data collected during picture viewing were analyzed, with eyes and lips as our areas of interests (AOIs). In both cases rectangles extending to sides of the face were used. For the eyes the, AOI spanned from the top of the eyebrows to an equivalent distance below the eyes, and for the lips the AOI extended to the bottom of the lips. For each AOI, time to first fixation, fixation time, and average fixation time were extracted. Four participants with tracking ratios below 80% were excluded from the analysis. Begaze was used to analyze the eye-movement data.

### Interpersonal reactivity index

Interpersonal reactivity index (IRI) was originally developed by Davis (1980, 1983a, 1983b). In this study we used the 16-item validated Persian version of IRI, which is used previously to investigate the general trait of empathy as well as its subcomponents (Golbabaei et al., 2023). Each item is responded on a 5-point Likert scale. The total score is in range of 0 and 64. The Questionnaire has shown a good internal consistency (.67-.71).

### Toronto Alexithymia scale

Toronto Alexithymia scale (TAS-20) is a 20-item questionnaire assessing three facets of alexithymia, namely difficulty in identifying feelings, difficulty in describing feelings, and externally oriented thinking. Each item is responded on a 5-point Likert scale. In this study we used a validated Farsi version (Besharat, 2008), which has been widely used previously and has shown a good internal consistency in range of .72 to .82.

### Autism-Spectrum Quotient

The 28-item shortened version of the Autism-Spectrum Quotient (AQ), developed by Hoekstra and colleagues (2010) as an abbreviated form of the original 50-item AQ by Baron-Cohen et al. (2001), assesses multiple domains related to autistic traits, in addition to providing an overall AQ score. Items are rated on a 4-point Likert scale ranging from definitely disagree to definitely agree. Higher scores reflect a stronger manifestation of autistic traits. The Farsi adaptation (Ashouri et al., 2020) used in the present study has demonstrated acceptable reliability of .61.

## Results

### Manipulation Check

To assess whether the empathy manipulation successfully differentiated the two conditions, we conducted paired t-tests comparing cognitive and affective empathy ratings across the high- and low-empathy conditions. Participants exhibited significantly greater cognitive empathy, *t*(69)=2.74, *p*=.008, as well as markedly higher affective empathy, *t*(69)=8.45, *p*<.001, in the high-empathy condition compared with the low-empathy condition (Figure-2).

**Figure 2.**
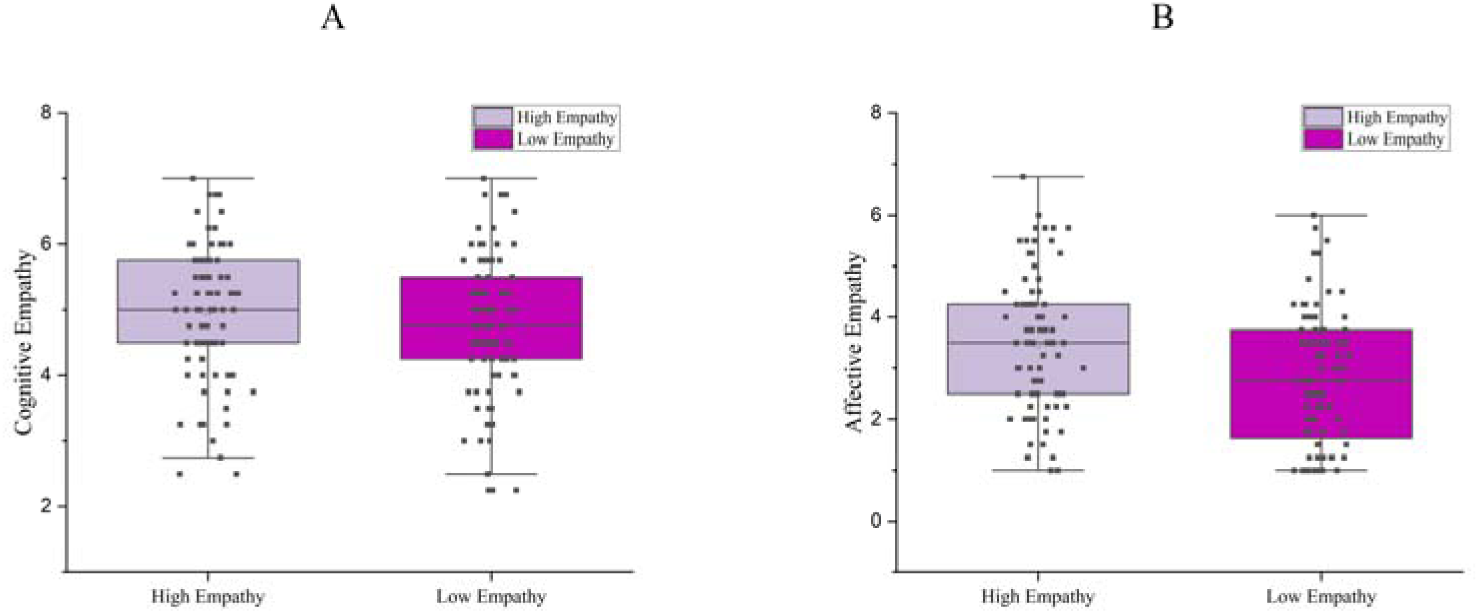
Comparison of Cognitive and Affective Empathy Across Conditions

### Time Perception

Using a one way repeated measure ANOVA we found a significant difference between three conditions in terms of time perception, *F*(2,168)=23.815, *p*<.001. Using a paired sample t-test we found the perception of time in high empathy condition to be significantly lower compared to both low empathy, *t*(69)=2.141, *p*=.036, and age estimation condition, *t*(69)=6.707, *p*<.001. Moreover, the perception of time in low empathy condition was also significantly lower than age estimation condition, *t*(69)=4.648, *p*<.001 (Figure-3). Since empathy manipulation was supposed to be the underlying reason for the difference in time perception among conditions, we next calculated the difference between the time perception in high vs low empathy conditions (from now on, called time perception_diff_; the lower the value, the lower the perceived duration in high empathy condition compared to the low empathy condition), and investigated the correlation between this difference and difference between cognitive empathy in both conditions, as well as the difference between the affective empathy in both conditions. We found a significant negative correlation between time perception_diff_ and both cognitive, *r*(69)=-.313, *p*=.008, and affective empathy, *r*(69)=-.304, *p*=.010 (Figure-4).

**Figure 3.**
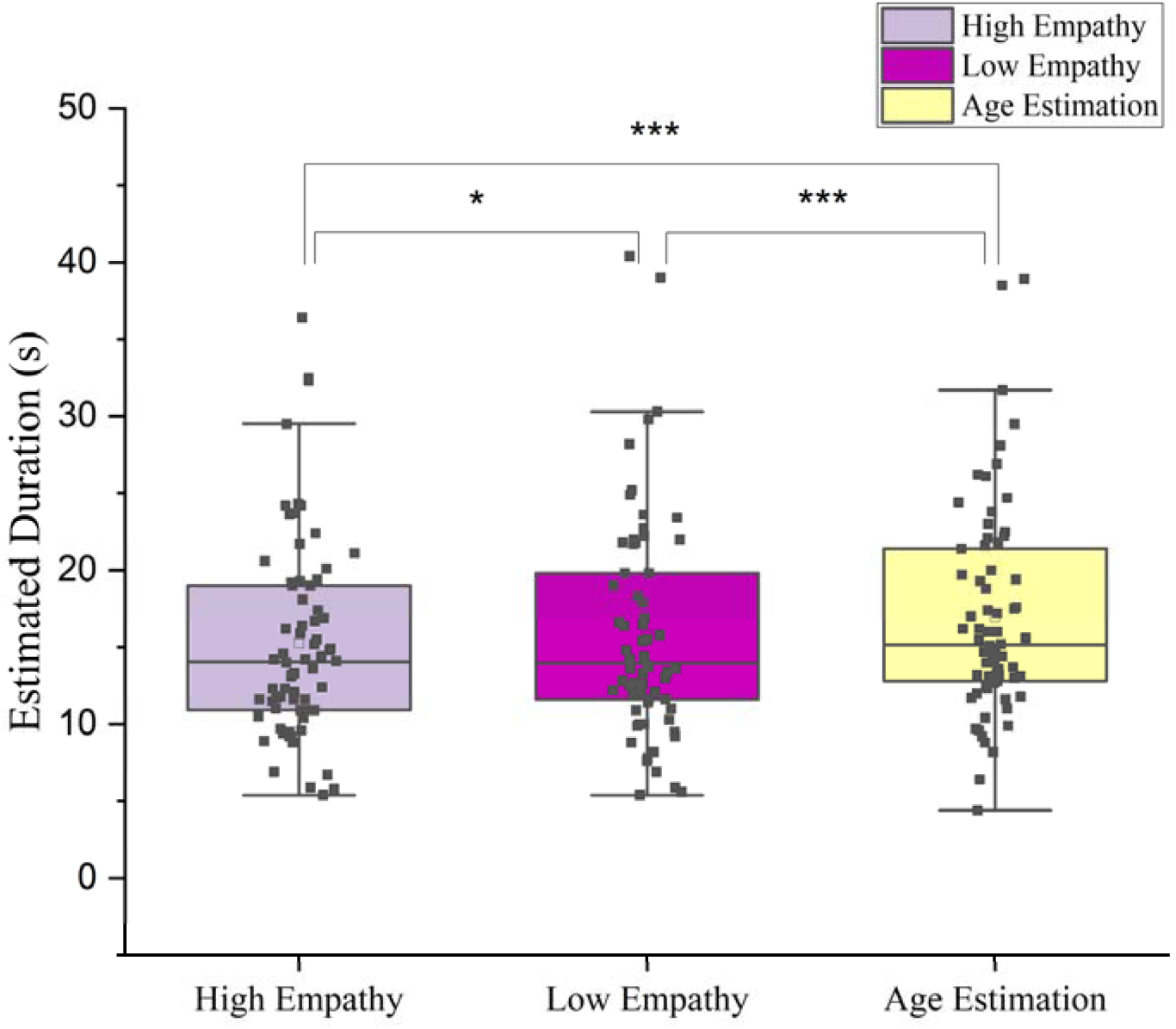
Estimated Duration Across Experimental Conditions

**Figure 4.**
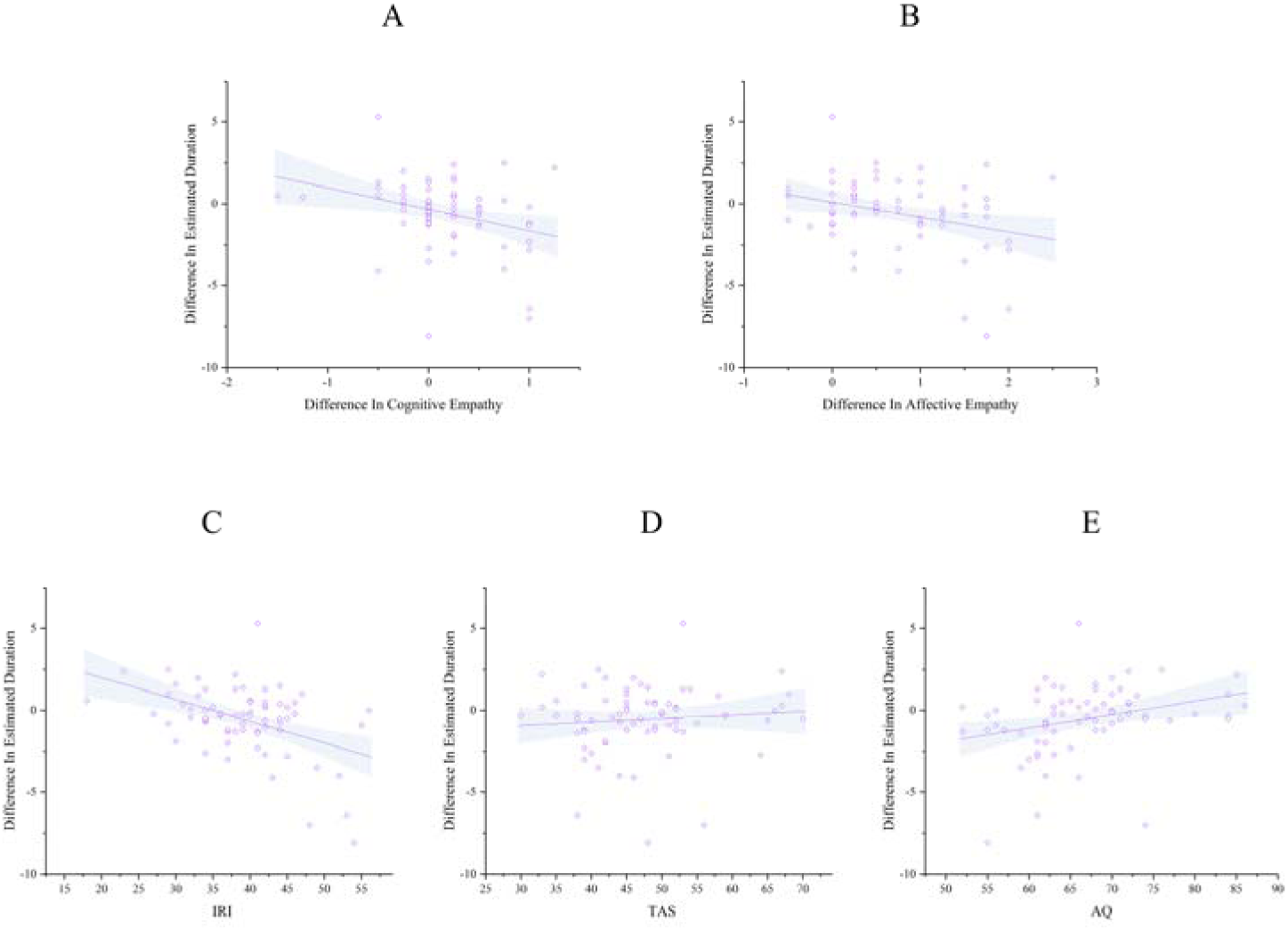
Associations Between Differences in Estimated Duration, State Empathy, and Personality Characteristics. Note. A: correlation between differences in estimated duration and cognitive empathy; B: correlation between differences in estimated duration and affective empathy; C: correlation between differences in estimated duration and interpersonal reactivity index; D: correlation between differences in estimated duration and Toronto alexithymia scale ; E: correlation between differences in estimated duration and autism quotient.

### Physiological Measures

We first evaluated the difference between two conditions in terms of both heart-rate and skin conductance measures. Participants exhibited a higher heart rate, *t*(55)=2.071, *p*=.043, lower IBI, *t*(55)= –2.036, *p* =.047, and increased skin conductance responses, *t*(57) = 2.038, *p*=.046 in high empathy compared to low empathy condition (Table-1). Next, we examined whether differences in estimated duration were associated with physiological measures across conditions. However, no significant correlations were observed (*Ps*>.05; Table-2 and Figure-5).

**Table 1.** Comparison of Physiological Arousal in High and Low Empathy Conditions.

| Measure | High Empathy |  | Low Empathy |  | t | p |
| --- | --- | --- | --- | --- | --- | --- |
|  | M | SD | M | SD |  |  |
| Heart Rate | 86.034 | 10.969 | 85.627 | 10.845 | 2.071 | .043 |
| IBI | 0.710 | 0.095 | 0.713 | 0.095 | -2.036 | .047 |
| SCR | 1.016 | 1.000 | 0.895 | .923 | 2.038 | .046 |

Notes. IBI: Interbeat Interval; SCR: Skin Conductance Response

**Table 2.** Correlation Between Difference in Estimated Duration and Physiological Arousal.

| Measure | M | SD | r | p |
| --- | --- | --- | --- | --- |
| Heart Rate<sub>diff</sub> | 0.407 | 1.469 | .028 | 0.839 |
| IBI<sub>diff</sub> | -0.003 | 0.013 | .017 | 0.903 |
| SCR<sub>diff</sub> | 0.121 | 0.451 | 0.037 | 0.781 |

Notes. Heart Rate<sub>diff</sub>: Difference in Heart Rate between High and Low Empathy conditions; IBI<sub>diff</sub>: Difference in Interbeat Interval between High and Low Empathy conditions; SCR<sub>diff</sub>: Difference in Skin Conductance Response between High and Low Empathy conditions;

**Figure 5.**
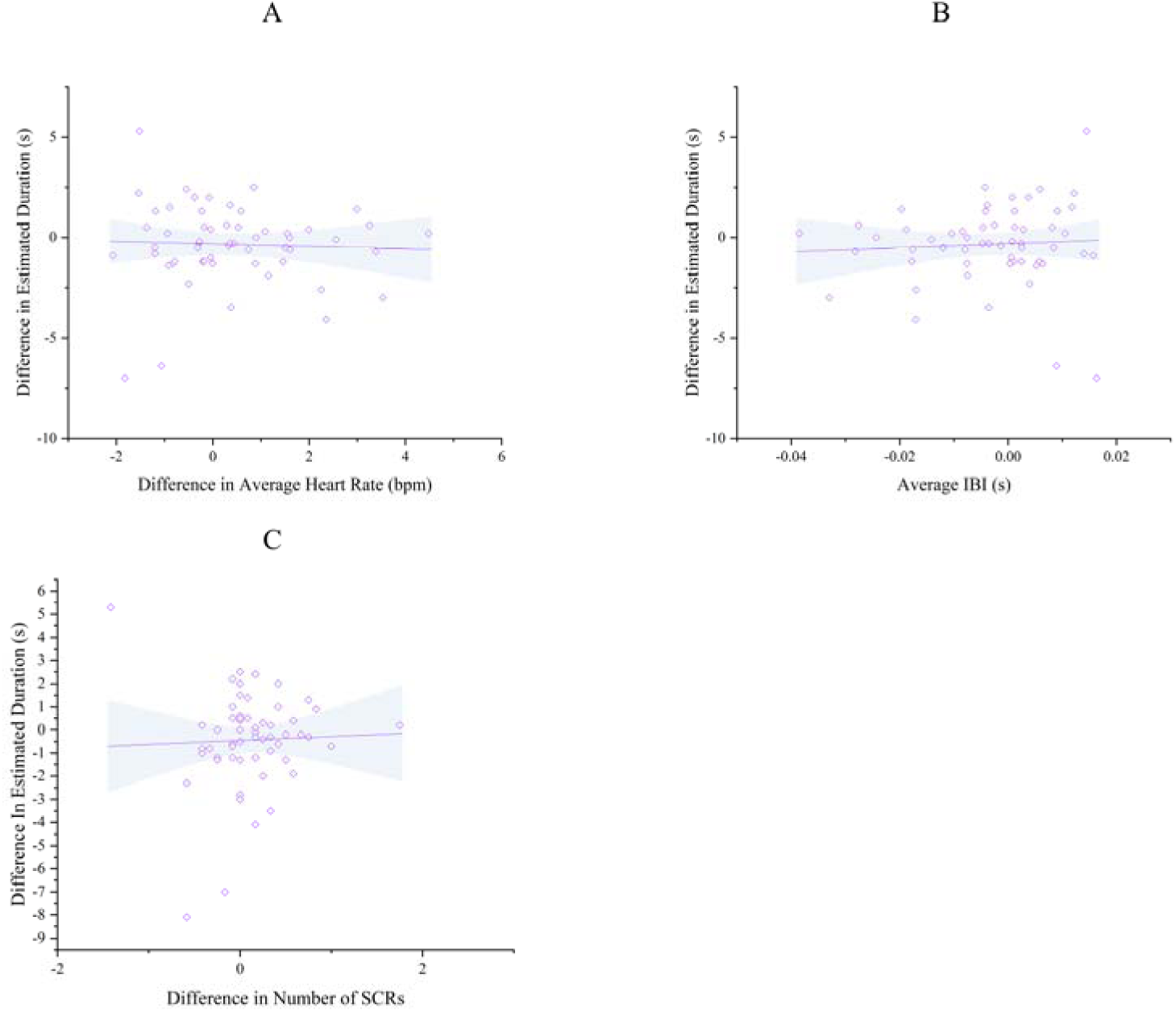
Associations Between Differences in Estimated Duration, Heart Rates, Interbeat Intervals, and Number of Skin Conductance Responses. Note. A: correlation between differences in estimated duration and average heart rate; B: correlation between differences in estimated duration and interbeat interval; C: correlation between differences in estimated duration and skin conductance response;

### Time perception and Eye-Movements

We first evaluated the difference between high and low empathy conditions in terms of eye-movement measures, among which we found a significant higher total, *t*(65)=3.865, *p*<.001, and average fixation duration, *t*(65)=2.570, *p*=.012, in the eye region in high empathy condition compared to low empathy condition (Table-3).

**Table 3.** Comparison of Eye-Movements in High and Low Empathy Conditions.

| Measure | High Empathy |  | Low Empathy |  | *t* | p |
| --- | --- | --- | --- | --- | --- | --- |
|  | *M* | *SD* | *M* | *SD* |  |  |
| Eyes |  |  |  |  |  |  |
| First Fixation Duration | 344.317 | 178.253 | 348.502 | 198.273 | -0.158 | 0.875 |
| Total Fixation Duration | 4149.775 | 2226.201 | 3625.713 | 1942.562 | 3.865 | 0.001 |
| Average Fixation Duration | 453.497 | 251.029 | 400.499 | 175.308 | 2.570 | 0.012 |
| Lips |  |  |  |  |  |  |
| First Fixation Duration | 368.134 | 201.462 | 398.398 | 209.378 | -1.248 | 0.216 |
| Total Fixation Duration | 1714.510 | 1057.595 | 1687.608 | 1007.465 | 0.379 | 0.706 |
| Average Fixation Duration | 389.877 | 195.582 | 402.833 | 201.441 | -0.640 | 0.525 |

We next evaluated the correlation between the difference in precepted time between conditions (Time perception_diff_ ) and difference between three measures of eye-movement in each condition for two areas of interest of eyes and lips. We found a significant negative different between Time perception_diff_ and First Fixation Duration_diff,_, *r*(65)= -.346, *p*=.004, and Average Fixation Duration_diff_, *r*(65)= -.246, *p*=.046, showing that longer the first fixation and the more the average fixation on eye region in the high empathy condition compared to low empathy condition, the shorter the perception of time in this condition (Table-4 and Figure-6).

**Table 4.** Correlation Between Difference in Estimated Duration and Eye-Movements.

| Measure | M | SD | r | p |
| --- | --- | --- | --- | --- |
| Eyes |  |  |  |  |
| First Fixation Duration<sub>diff</sub> | -4.185 | 215.222 | -.346 | .004 |
| Total Fixation Duration<sub>diff</sub> | 524.062 | 1101.586 | -.181 | .146 |
| Average Fixation Duration<sub>diff</sub> | -2.778 | 171.371 | -.246 | .046 |
| Lips |  |  |  |  |
| First Fixation Duration<sub>diff</sub> | -30.265 | 196.996 | .053 | .672 |
| Total Fixation Duration<sub>diff</sub> | 26.903 | 576.381 | -.084 | .502 |
| Average Fixation Duration<sub>diff</sub> | -12.956 | 164.523 | .076 | .545 |

**Figure 6.**
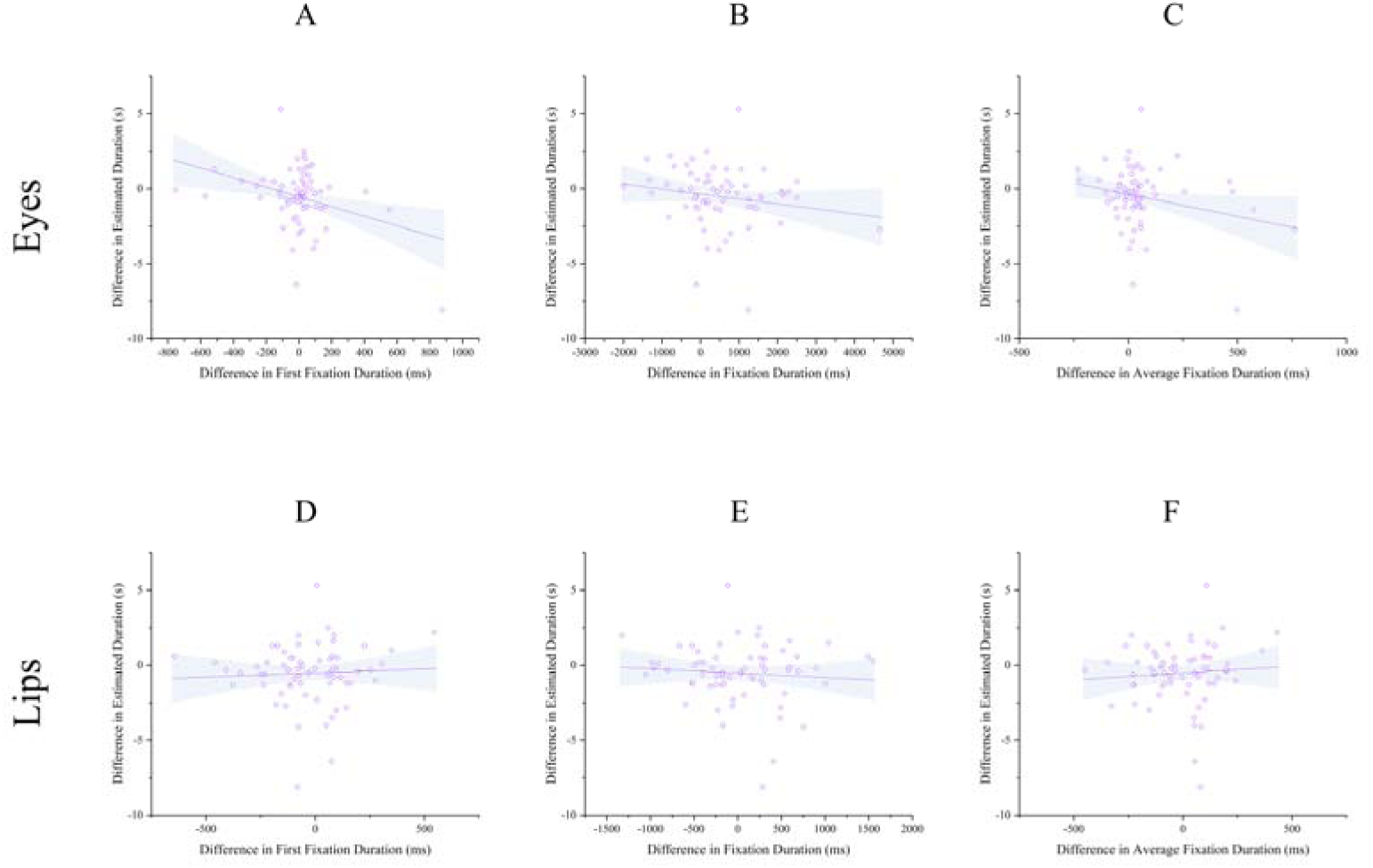
Associations Between Differences in Estimated Duration and Eye-Movements. Note. A: correlation between differences in estimated duration and first fixation duration in the eye region; B: correlation between differences in estimated duration and fixation duration in the eye region; C: correlation between differences in estimated duration and average fixation duration in the eye region; D: correlation between differences in estimated duration and first fixation duration in the lips region; E: correlation between differences in estimated duration and fixation duration in the lips region; C: correlation between differences in estimated duration and average fixation duration in the lips region;

### Time Perception and Personality Traits

We next evaluated the correlation between the time perception_diff_ and scores derived from three questionnaires of IRI, TAS-20, and AQ. Time perception_diff_ showed a significant negative correlation with IRI and, *r*(69)=-.452, *p*<.001, and a significant positive correlation with AQ, *r*(69)=.309, *p*=.009, but no relationship with TAS-20, *r*(69)=.090, *p* =.459 (Figure-4).

## Discussion

Considering the complex nature of empathic phenomenon, the present study examined how empathy alters time perception. While the early accounts of internal clock propose that emotional arousal lengthens subjective duration, frameworks grounded in motivational dimensional model, shared representations of empathy and self-construal theory predict the opposite pattern. Our results support this latter perspective showing that empathizing with another person shortens perceived time. Rather than contradicting internal clock models, these findings highlight their complexity and contribute to explaining inconsistencies in the literature.

A key insight emerging from our result is that empathy appears to reduce perceived temporal distance in a manner analogous to its effects on psychological distance and more broadly. Empathy is consistently associated with reduced self-other distinction (Cooke et al., 2018; Lamm et al., 2016), and decreased perception of distance, both at psychological (Stupacher et al., 2022) and physical (Golbabaei & Borhani, 2024) levels. Prior work suggests that reductions in one level of construal, such as dimension of psychological distance tend to generalize to others, including temporal distance (Fiedler et al., 2012; Kanten, 2011; Rim et al., 2009; Trope & Liberman, 2010). Besides, neural underlying of time perception and spatial perception are also highly overlapped (Robbe, 2021; Unruh et al., 2023). In line with this, shorter duration estimation due to enhanced self-other overlap has also been reported in embodiment paradigms such as the rubber-hand illusion and enfacement (Suzuki et al., 2019; Unruh et al., 2023). The proposed mechanism is that greater congruency and plausibility between self and other reduce prediction error, resulting in a subjectively faster passage of time (Unruh et al., 2021, 2023). During empathic engagement, individuals often vicariously experience others’ feelings, a process akin to the multisensory integration underlying enfacement or whole-body illusions. The incorporation of another’s affective state into one’s own bodily experience may accelerate the subjective passage of time. By enhancing self–other overlap and diminishing self–other distinction, empathy may therefore foster a comparable compression of temporal experience.

Motivational mechanisms may also contribute. According to motivational dimensional model of time perception (Gable et al., 2016, 2022; Gable & Poole, 2012), approach motivation accelerates the perceived passage of time, whereas avoidance slows it (Gable et al., 2016; Gable & Poole, 2012). Shorter perceived time has been proposed to enhance approach motivation, especially when accompanied by a burden or negative affect. Under these conditions, individuals are more inclined to remove the obstacle or pursue the goal, and this prolonged engagement is adaptive for problem solving (Gable et al., 2016). Moreover, feeling closer to something in terms of time or distance leads to more motivation to act and lower self-other distinction (Wohl & McGrath, 2007). Apart from sharing others’ emotions and understanding their feelings, empathy is intrinsically linked to approach motivation, orienting individuals toward others’ needs and helping to alleviate their sufferings (Eisenberg et al., 2010; Kozakevich Arbel et al., 2024; Samani et al., 2022; Zaki, 2020). Thus, temporal contraction during empathic engagement may serve an adaptive function, energizing individuals to act, especially when the target is suffering. Consistent with this idea, previous work has shown that how people perceive time influences their willingness to help (Latoschik & Wienrich, 2022; Niiya et al., 2025; Unruh et al., 2021). Research on procrastination further suggests that when time feels scarce or compressed, individuals perceive tasks as more urgent and are more motivated to act. Empathic time compression may therefore support prosocial responding by creating a sense of immediacy.

Another important finding was that both high and low empathy conditions led to shorter duration estimates, compared to the age estimation condition, which imposes attentional demands but lacked empathic involvement. Although attentional load and diversion are known to influence timing (Dunn et al., 2017; Im & Varma, 2018; Tamm et al., 2014), our results suggest that the observed empathy effects cannot be attributed to differences in attentional resources.

Critically, both cognitive and affective empathy scores were negatively associated with changes in time perception, where the higher the cognitive or affective empathy, the lower the perceived time. This is of great importance, and reinforces that the effects observed here systematically related to empathic tendencies and are not artifacts of the task. It further indicates that individual differences in empathy are also traceable, and shape fundamental perceptual processes, including subjective time.

Moreover, in line with our expectations high empathy trials elicited increased heart rate, reduced interbeat interval, and elevated skin conductance response. This is in line with previous studies showing that empathy is related to increased arousal or changes in physiological signals (Deuter et al., 2018; Golbabaei, et al., 2022; Hasan, Hossain, Ghosh, et al., 2025; Hasan, Hossain, Krishna, et al., 2025). However, these physiological changes did not predict alterations in time perception. This finding challenges the view that arousal serves as the primary internal pulse driving the internal clock, suggesting instead that it is not the sole source for the hypothetical accumulator and that additional factors or signals must also contribute. Moreover, temporal compression during empathy is not solely driven by autonomic activation. Besides, it aligns with theoretical accounts such as motivated dimensional model of time perception emphasizing that arousal and motivation should not be conflated. Arousal during approach-motivated states does not necessarily lengthen time and may even accompany perceived time acceleration or have no effect (Droit-Volet & Gil, 2016; Gable et al., 2022). On the other hand, while physiological signals play a role in empathy, social-cognitive dimensions such as self–other distinction, relevant for time-perception changes, may depend less on isolated physiological responses and more on the synchrony between the empathizer’s and the target’s physiological and behavioral states (Ebrahimi et al., 2024; Goldstein et al., 2017; Qaiser et al., 2023; but also see Pan et al., 2023). Thus, it is possible that the critical factor underlying changes in time perception is not the physiological response of a single individual, but the interaction or coupling of physiological signals between both sides of the empathic process.

Visual attention also provided informative insights. First, we showed that greater visual attention to eyes is related to empathic responding. This aligns with previous work demonstrating that the eyes are among the most salient and informative cues for detecting pain, and that increased gaze to the eyes corresponds to heightened empathy (Cowan et al., 2014; Golbabaei, et al., 2023; Golbabaei & Borhani, 2024). More importantly, we found that both longer initial fixations and greater average fixation time on the eyes predicted a faster passage of time. Previous work indicates that such eye-focused attention predicts changes in perceived distance during empathy. Our findings extend this effect to temporal perception. One speculative interpretation is that enhanced attention to the eyes increases access to socially relevant visual cues, which must be integrated with somatosensory resonance processes. This integration may strengthen shared representations and self–other overlap, contributing to compressed temporal experience. Since we did not directly measure self–other overlap, this explanation remains tentative.

Finally, in line with our expectation, trait-level variables, namely trait empathy and autism-spectrum quotient, also modulated empathy-related changes in temporal perception. Trait and state empathy are distinct but related constructs (Cuff et al., 2014). Dispositional empathy has been linked to greater self–other overlap (Bukowski et al., 2020), and individuals high in trait empathy typically show stronger motivation to approach and alleviate others’ pain (Molnar-Szakacs, 2011; Pavey et al., 2012; Samani et al., 2022; Zaki, 2014). Conversely, higher autism-spectrum traits were associated with reduced temporal contraction. Autism is characterized by difficulties in empathy (Adler et al., 2015; Shamay-Tsoory et al., 2002) and atypical self–other processing (Hoffmann et al., 2015; Lavenne-Collot et al., 2023). Our results suggest that these characteristics extend to the perceptual consequences of empathic phenomenon. Notably, prior work has documented alterations in general time perception among ASD individuals, but these findings are inconsistent and largely confined to higher-order temporal judgments rather than duration estimation (Casassus et al., 2019; Vatakis et al., 2015; Wallace & Happé, 2008). Our findings indicate that autistic traits modulate how empathy reshapes temporal processing, rather than time perception in isolation.

In contrast to our expectation, alexithymia did not modulate empathy-related changes in time perception. Although alexithymia is strongly associated with empathy difficulties, its relationship with temporal perception remains largely unexplored. One study suggests alexithymia may lead to overestimated durations for negative stimuli (Vicario et al., 2023), but evidence is scarce. It is possible that alexithymia entails its own time perception abnormalities, driven by mechanisms independent of empathy, thereby confounding its relationship with empathy-induced temporal changes. This highlights the need for future research that disentangles these effects.

Taken together, our findings demonstrate that empathy shortens subjective time, likely through shared bodily representations and approach motivation rather than arousal alone. This finding enriches internal clock framework by demonstrating that arousal does not invariably lead to temporal expansion, highlighting the role of more complex cognitive and affective processes in temporal judgment. They also emphasize the central role of visual attention, particularly to the eyes as a highly informative source, in shaping empathic and temporal processes. Finally, we show that trait empathy facilitates, and autism-spectrum traits attenuate, these perceptual consequences of empathy.

Yet, our study also has limitations. We did not directly measure self-other distinction and motivation to help, which restricts the strength of inferences regarding these mechanisms. Future work could incorporate established self–other distinction measures or manipulate approach motivation experimentally. In addition, empathy was examined in a single-person design, preventing investigation of the role of physiological synchrony in modulating time perception. Moreover, to allow manipulation of empathy and reliable physiological recording, stimuli were not presented across a wide range of durations, including very short and very long intervals. Finally, since alexithymia shares substantial variance with depression, which itself influences time perception, future studies should statistically control for depression to isolate the unique contribution of alexithymia.

## Data Availability

The data that support the findings of this study are available upon reasonable request from the corresponding author.

## Conflicts of Interest

Authors have no conflict of interest to declare.

## Funding

No external funding was received for the conduct of this study.

## Authors Contribution

SG: Conceptualization; Data Collection; Formal analysis; Investigation; Methodology; Resources; Software; Visualization; Writing—Original draft; Writing—Review & editing. MH: Data Collection; Writing—Review & editing. ZK: Data Collection; Writing—Review & editing. KB: Conceptualization; Formal analysis; Methodology; Project administration; Resources; Supervision; Writing—Original draft; Writing—Review & editing.

## Notes

### Competing Interest Statement

The authors have declared no competing interest.

## References

Adler, N., Dvash, J., & Shamay-Tsoory, S. G. (2015). Empathic Embarrassment Accuracy in Autism Spectrum Disorder. Autism Research, 8(3), 241–249. 10.1002/AUR.1439;REQUESTEDJOURNAL:JOURNAL:19393806;WGROUP:STRING:PUBLICATION

Ashouri, A., Asgharzade, A., Ebrahimi, A., Ghojazadeh, M., & Akbarzadeh, A. (2020). Psychometric Properties of Persian Version of Autism-Spectrum Quotient (AQP-28): Evidence from Iranian Non-clinical Sample. 10.21203/rs.3.rs-47383/v1

Bagby, R. M., Parker, J. D. A., & Taylor, G. J. (1994). The twenty-item Toronto Alexithymia scale-I. Item selection and cross-validation of the factor structure. Journal of Psychosomatic Research. 10.1016/0022-3999(94)90005-1

Baron-Cohen, S., Wheelwright, S., Skinner, R., Martin, J., & Clubley, E. (2001). The Autism-Spectrum Quotient (AQ): Evidence from Asperger Syndrome/High-Functioning Autism, Males and Females, Scientists and Mathematicians. Journal of Autism and Developmental Disorders, 31(1), 5–17. 10.1023/A:1005653411471

Benau, E. M., & Atchley, R. A. (2020). Time flies faster when you’re feeling blue: sad mood induction accelerates the perception of time in a temporal judgment task. Cognitive Processing *2020 21:3*, 21(3), 479–491. 10.1007/S10339-020-00966-8

Besharat, M. A. (2008). Assessing Reliability and Validity of the Farsi Version of the Toronto Alexithymia Scale in a Sample of Substance-Using Patients. 10.2466/PR0.102.1.259-270

Bilali, R., Iqbal, Y., & Çelik, A. B. (2018). The role of national identity, religious identity, and intergroup contact on social distance across multiple social divides in Turkey. International Journal of Intercultural Relations, 65, 73–85. 10.1016/J.IJINTREL.2018.04.007

Bird, G., & Viding, E. (2014). The self to other model of empathy: Providing a new framework for understanding empathy impairments in psychopathy, autism, and alexithymia. Neuroscience & Biobehavioral Reviews, 47, 520–532. 10.1016/J.NEUBIOREV.2014.09.021

Björkqvist, K., Österman, K., & Kaukiainen, A. (2000). Social intelligence − empathy = aggression? Aggression and Violent Behavior, 5(2), 191–200. 10.1016/S1359-1789(98)00029-9

Bukowski, H., Tik, M., Silani, G., Ruff, C. C., Windischberger, C., & Lamm, C. (2020). When differences matter: rTMS/fMRI reveals how differences in dispositional empathy translate to distinct neural underpinnings of self-other distinction in empathy. Cortex, 128, 143–161. 10.1016/J.CORTEX.2020.03.009

Casassus, M., Poliakoff, E., Gowen, E., Poole, D., & Jones, L. A. (2019). Time perception and autistic spectrum condition: A systematic review. Autism Research, 12(10), 1440–1462. 10.1002/AUR.2170;PAGE:STRING:ARTICLE/CHAPTER

Chambon, M., Droit-Volet, S., & Niedenthal, P. M. (2008). The effect of embodying the elderly on time perception. Journal of Experimental Social Psychology, , 44(3), 672–678. 10.1016/j.jesp.2007.04.014

Colonnello, V., Domes, G., & Heinrichs, M. (2016). As time goes by: Oxytocin influences the subjective perception of time in a social context. Psychoneuroendocrinology, 68, 69–73. 10.1016/J.PSYNEUEN.2016.02.015

Cooke, A. N., Bazzini, D. G., Curtin, L. A., & Emery, L. J. (2018). Empathic understanding: Benefits of perspective-taking and facial mimicry instructions are mediated by self-other overlap. Motivation and Emotion *2018 42:3*, 42(3), 446–457. 10.1007/S11031-018-9671-9

Cowan, D. G., Vanman, E. J., & Nielsen, M. (2014). Motivated empathy: The mechanics of the empathic gaze. Cognition and Emotion. 10.1080/02699931.2014.890563

Cuff, B. M. P., Brown, S. J., Taylor, L., & Howat, D. J. (2014). Empathy: A review of the concept. In Emotion Review. 10.1177/1754073914558466

Davis, M. H. (1983a). Measuring individual differences in empathy: Evidence for a multidimensional approach. Journal of Personality and Social Psychology, 44(1), 113–126. 10.1037/0022-3514.44.1.113

Davis, M. H. (1983b). The effects of dispositional empathy on emotional reactions and helping: A multidimensional approach. Journal of personality, 51(2), 167–184.

Davis, M. H., & Association, A. P. (1980). A multidimensional approach to individual differences in empathy. JSAS Catalog of Selected Documents in Psychology.

Decety, J., & Cowell, J. M. (2014). The complex relation between morality and empathy. In Trends in Cognitive Sciences. 10.1016/j.tics.2014.04.008

Deuter, C. E., Nowacki, J., Wingenfeld, K., Kuehl, L. K., Finke, J. B., Dziobek, I., & Otte, C. (2018). The role of physiological arousal for self-reported emotional empathy. Autonomic Neuroscience: Basic and Clinical, 214, 9–14. 10.1016/j.autneu.2018.07.002

Dirnberger, G., Hesselmann, G., Roiser, J. P., Preminger, S., Jahanshahi, M., & Paz, R. (2012). Give it time: Neural evidence for distorted time perception and enhanced memory encoding in emotional situations. NeuroImage, 63(1), 591–599. 10.1016/J.NEUROIMAGE.2012.06.041

Drew, M. R., Fairhurst, S., Malapani, C., Horvitz, J. C., & Balsam, P. D. (2003). Effects of dopamine antagonists on the timing of two intervals. Pharmacology Biochemistry and Behavior, 75(1), 9–15. 10.1016/S0091-3057(03)00036-4

Droit-Volet, S., Brunot, S., & Niedenthal, P. M. (2004). Perception of the duration of emotional events. Cognition and Emotion, 18(6), 849–858. 10.1080/02699930341000194;WEBSITE:WEBSITE:TFOPB;PAGEGROUP:STRING:PUBLICATION

Droit-Volet, S., Fayolle, S. L., & Gil, S. (2011). Emotion and time perception: Effects of film-induced mood. Frontiers in Integrative Neuroscience, 5, 11974. 10.3389/FNINT.2011.00033/BIBTEX

Droit-Volet, S., & Gil, S. (2016). The emotional body and time perception. Cognition and Emotion, 30(4), 687–699. 10.1080/02699931.2015.1023180;SUBPAGE:STRING:FULL

Droit-Volet, S., & Wearden, J. (2002). Speeding up an internal clock in children? Effects of visual flicker on subjective duration. Quarterly Journal of Experimental Psychology Section B: Comparative and Physiological Psychology, 55(3), 193–211. 10.1080/02724990143000252;SUBPAGE:STRING:ABSTRACT;WGROUP:STRING:PUBLICATION

Dunn, T. L., Inzlicht, M., & Risko, E. F. (2017). Anticipating cognitive effort: roles of perceived error-likelihood and time demands. Psychological Research *2017 83:5*, 83(5), 1033–1056. 10.1007/S00426-017-0943-X

Dvash, J., & Shamay-Tsoory, S. G. (2014). Theory of mind and empathy as multidimensional constructs: Neurological foundations. Topics in Language Disorders, 34(4), 282–295. 10.1097/TLD.0000000000000040

Ebrahimi, F., Borhani, K., & Vahabie, A. (2024). The Effect of Behavioral Synchrony on Choosing Compensation in Adolescents and Adults’ Decision-Making. Journal of Applied Psychological Research. 10.22059/JAPR.2024.361306.644661

Eisenberg, N., Eggum, N. D., & Di Giunta, L. (2010). Empathy-Related Responding: Associations with Prosocial Behavior, Aggression, and Intergroup Relations. Social Issues and Policy Review, 4(1), 143–180. 10.1111/j.1751-2409.2010.01020.x

Farmer, H., & Maister, L. (2017). Putting Ourselves in Another’s Skin: Using the Plasticity of Self-Perception to Enhance Empathy and Decrease Prejudice. Social Justice Research *2017 30:4*, 30(4), 323–354. 10.1007/S11211-017-0294-1

Faul, F., Erdfelder, E., Lang, A. G., & Buchner, A. (2007). G*Power 3: A flexible statistical power analysis program for the social, behavioral, and biomedical sciences. Behavior Research Methods, 39(2), 175–191. 10.3758/BF03193146/METRICS

Fiedler, K., Jung, J., Wänke, M., & Alexopoulos, T. (2012). On the relations between distinct aspects of psychological distance: An ecological basis of construal-level theory. Journal of Experimental Social Psychology, 48(5), 1014–1021. 10.1016/j.jesp.2012.03.013

Gable, P. A., Neal, L. B., & Poole, B. D. (2016). Sadness speeds and disgust drags: Influence of motivational direction on time perception in negative affect. Motivation Science, 2(4), 238–255. 10.1037/MOT0000044

Gable, P. A., & Poole, B. D. (2012). Time Flies When You’re Having Approach-Motivated Fun: Effects of Motivational Intensity on Time Perception. Psychological Science, 23(8), 879–886. 10.1177/0956797611435817

Gable, P. A., Wilhelm, A. L., & Poole, B. D. (2022). How Does Emotion Influence Time Perception? A Review of Evidence Linking Emotional Motivation and Time Processing. Frontiers in Psychology, 13, 848154. 10.3389/FPSYG.2022.848154/BIBTEX

Gibbon, J. (1977). Scalar expectancy theory and Weber’s law in animal timing. Psychological Review, 84(3), 279–325. 10.1037/0033-295X.84.3.279

Gibbon, J., Church, R. M., & Meck, W. H. (1984). Scalar Timing in Memory. Annals of the New York Academy of Sciences, 423(1), 52–77. 10.1111/J.1749-6632.1984.TB23417.X

Gibbons, S. B. (2011). Understanding Empathy as a Complex Construct: A Review of the Literature. In Clinical Social Work Journal. 10.1007/s10615-010-0305-2

Gil, S., & Droit-Volet, S. (2011). Time perception in response to ashamed faces in children and adults. Scandinavian Journal of Psychology, 52(2), 138–145. 10.1111/J.1467-9450.2010.00858.X

Golbabaei, S., Barati, M., Haromi, M. E., Ghazazani, N., & Borhani, K. (2022). Development and construct validation of a short form of the interpersonal reactivity index in Iranian community. Current Psychology, 11(4), 217–228. 10.1007/s12144-022-02716-9

Golbabaei, S., & Borhani, K. (2024). Nearsighted empathy: exploring the effect of empathy on distance perception, with eye movements as modulators. Scientific Reports, 14(1). 10.1038/S41598-024-76731-0

Golbabaei, S., Sammaknejad, N., & Borhani, K. (2022). Physiological Indicators of The Relation Between Autistic Traits and Empathy: Evidence From Electrocardiogram and Skin Conductance Signals. 2022 29th National and 7th International Iranian Conference on Biomedical Engineering, ICBME 2022, 177–183. 10.1109/ICBME57741.2022.10053068

Golbabaei, S., Sammaknejad, N., & Borhani, K. (2023). Cognitive and Affective Empathy in Individuals with High and Low Alexithymia: The Mediating Role of Eye-Gaze Pattern to Facial Expressions. Quarterly of Applied Psychology, 17(3), 51–75. 10.48308/APSY.2023.230817.1460

Goldstein, P., Weissman-Fogel, I., & Shamay-Tsoory, S. G. (2017). The role of touch in regulating inter-partner physiological coupling during empathy for pain. Scientific Reports. 10.1038/s41598-017-03627-7

Grynberg, D., Luminet, O., Corneille, O., Grèzes, J., & Berthoz, S. (2010). Alexithymia in the interpersonal domain: A general deficit of empathy? Personality and Individual Differences, 49(8), 845–850. 10.1016/j.paid.2010.07.013

Hall, J. A., & Schwartz, R. (2019). Empathy present and future. The Journal of Social Psychology, 159(3), 225–243. 10.1080/00224545.2018.1477442

Hasan, M. R., Hossain, M. Z., Ghosh, S., Krishna, A., & Gedeon, T. (2025). Empathy Detection from Text, Audiovisual, Audio or Physiological Signals: A Systematic Review of Task Formulations and Machine Learning Methods. IEEE Transactions on Affective Computing. 10.1109/TAFFC.2025.3590107

Hasan, M. R., Hossain, M. Z., Krishna, A., Rahman, S., & Gedeon, T. (2025). Are You Really Empathic? Evidence from Trait, State and Speaker-Perceived Empathy, and Physiological Signals. 1. https://arxiv.org/pdf/2509.16923

Hoekstra, R. A., Vinkhuyzen, A. A. E., Wheelwright, S., Bartels, M., Boomsma, D. I., Baron-Cohen, S., Posthuma, D., & Van Der Sluis, S. (2010). The Construction and Validation of an Abridged Version of the Autism-Spectrum Quotient (AQ-Short). Journal of Autism and Developmental Disorders *2010 41:5*, 41(5), 589–596. 10.1007/S10803-010-1073-0

Hoffmann, F., Koehne, S., Steinbeis, N., Dziobek, I., & Singer, T. (2015). Preserved Self-other Distinction During Empathy in Autism is Linked to Network Integrity of Right Supramarginal Gyrus. Journal of Autism and Developmental Disorders *2015 46:2*, 46(2), 637–648. 10.1007/S10803-015-2609-0

Im, S. hyun, & Varma, S. (2018). Distorted Time Perception during Flow as Revealed by an Attention-Demanding Cognitive Task. Creativity Research Journal, 30(3), 295–304. 10.1080/10400419.2018.1488346

Kanten, A. B. (2011). The effect of construal level on predictions of task duration. Journal of Experimental Social Psychology, 47(6), 1037–1047. 10.1016/J.JESP.2011.04.005

Kliegl, K. M., Limbrecht-Ecklundt, K., Dürr, L., Traue, H. C., & Huckauf, A. (2015). The complex duration perception of emotional faces: Effects of face direction. Frontiers in Psychology, 6(MAR), 128147. 10.3389/FPSYG.2015.00262/BIBTEX

Kozakevich Arbel, E., Shamay-Tsoory, S. G., & Hertz, U. (2024). Adaptive empathic response selection is sensitive to multiple dimensions of social interaction. Communications Psychology *2024 2:1*, 2(1), 112-. 10.1038/s44271-024-00164-8

Lamm, C., Bukowski, H., & Silani, G. (2016). From shared to distinct self-other representations in empathy: Evidence from neurotypical function and socio-cognitive disorders. Philosophical Transactions of the Royal Society B: Biological Sciences, 371(1686). 10.1098/rstb.2015.0083

Latoschik, M. E., & Wienrich, C. (2022). Congruence and Plausibility, Not Presence: Pivotal Conditions for XR Experiences and Effects, a Novel Approach. Frontiers in Virtual Reality, 3, 694433. 10.3389/FRVIR.2022.694433/BIBTEX

Lavenne-Collot, N., Tersiguel, M., Dissaux, N., Degrez, C., Bronsard, G., Botbol, M., & Berthoz, A. (2023). Self/other distinction in adolescents with autism spectrum disorder (ASD) assessed with a double mirror paradigm. PLOS ONE, 18(3), e0275018. 10.1371/JOURNAL.PONE.0275018

Li, W., He, Q. F., Lan, J. Z., Attiq-Ur-Rehman, Ge, M. W., Shen, L. T., Hu, F. H., Jia, Y. J., & Chen, H. L. (2024). Empathy as a Mediator of the Relation between Peer Influence and Prosocial Behavior in Adolescence: A Meta-Analysis. Journal of Youth and Adolescence *2024 54:3*, 54(3), 682–703. 10.1007/S10964-024-02079-3

Lopez, C., Falconer, C. J., & Mast, F. W. (2013). Being Moved by the Self and Others: Influence of Empathy on Self-Motion Perception. PLOS ONE, 8(1), e48293. 10.1371/JOURNAL.PONE.0048293

Martínez-Pernía, D., Cea, I., Troncoso, A., Blanco, K., Calderón Vergara, J., Baquedano, C., Araya-Veliz, C., Useros-Olmo, A., Huepe, D., Carrera, V., Mack Silva, V., & Vergara, M. (2023). “I am feeling tension in my whole body”: An experimental phenomenological study of empathy for pain. Frontiers in Psychology, 13, 999227. 10.3389/FPSYG.2022.999227/BIBTEX

Méndez Fernández, A. B., Lombardero Posada, X., Aguiar Fernández, F. X., Murcia Álvarez, E., & González Fernández, A. (2022). Professional preference for mental illness: The role of contact, empathy, and stigma in Spanish Social Work undergraduates. Health and Social Care in the Community, 30(4), 1492–1503. 10.1111/HSC.13479

Molnar-Szakacs, I. (2011). From actions to empathy and morality – A neural perspective. Journal of Economic Behavior & Organization, 77(1), 76–85. 10.1016/J.JEBO.2010.02.019

Moreira, D., Azeredo, A., Leite, Â., & Barbosa, F. (2025). Effects of impulsivity and emotions on time perception: Laboratory behavioral measures. Perception, 54(4), 239–251. 10.1177/03010066251316457

Morelli, S. A., Rameson, L. T., & Lieberman, M. D. (2014). The neural components of empathy: Predicting daily prosocial behavior. Social Cognitive and Affective Neuroscience, 9(1), 39–47. 10.1093/SCAN/NSS088

Niiya, Y., Yakin, S., Park, L. E., & Chang, Y. H. (2025). Nonzero-Sum Time Perception Is Associated with Greater Willingness to Help. European Journal of Investigation in Health, Psychology and Education, 15(5), 90. 10.3390/EJIHPE15050090/S1

Pan, H., Chen, Z., Jospe, K., Gao, Q., Sheng, J., Gao, Z., & Perry, A. (2023). Mood congruency affects physiological synchrony but not empathic accuracy in a naturalistic empathy task. Biological Psychology, 184, 108720. 10.1016/J.BIOPSYCHO.2023.108720

Pavey, L., Greitemeyer, T., & Sparks, P. (2012). “I help because i want to, not because you tell me to”: Empathy increases autonomously motivated helping. Personality and Social Psychology Bulletin, 38(5), 681–689. 10.1177/0146167211435940

Perry, A., Mankuta, D., & Shamay-Tsoory, S. G. (2015). OT promotes closer interpersonal distance among highly empathic individuals. Social Cognitive and Affective Neuroscience, 10(1), 3–9. 10.1093/SCAN/NSU017

Qaiser, J., Leonhardt, N. D., Le, B. M., Gordon, A. M., Impett, E. A., & Stellar, J. E. (2023). Shared Hearts and Minds: Physiological Synchrony During Empathy. Affective Science *2023 4:4*, 4(4), 711–721. 10.1007/S42761-023-00210-4

Reniers, R. L. E. P., Corcoran, R., Drake, R., Shryane, N. M., & Völlm, B. A. (2011). The QCAE: A questionnaire of cognitive and affective empathy. Journal of Personality Assessment. 10.1080/00223891.2010.528484

Rim, S. Y., Uleman, J. S., & Trope, Y. (2009). Spontaneous trait inference and construal level theory: Psychological distance increases nonconscious trait thinking. Journal of Experimental Social Psychology, 45(5), 1088–1097. 10.1016/J.JESP.2009.06.015

Robbe, D. (2021). Lost in time: rethinking duration estimation outside the brain. *(Preprint). PsyArXiv*. 10.31234/osf.io/3bcfy.

Samani, H., Golbabaei, S., & Borhani, K. (2022). Predicting Paternalism Based on Components of Empathy and Behavioral Contagion. Social Psychology Research, 12(46), 101–122. 10.22034/SPR.2022.333641.1737

Shamay-Tsoory, S. G., Tomer, R., Yaniv, S., & Aharon-Peretz, J. (2002). Empathy Deficits in Asperger Syndrome: a Cognitive Profile. Neurocase, 8(3), 245–252. 10.1093/NEUCAS/8.3.245

Singer, T., Dolan, R. J., & Frith, C. D. (2004). Empathy for Pain Involves the Affective but not Sensory Components of Pain. Science. 303 (5661): 1157–1162. 10.1126/science.1093535

Smith, A. (2017). The Empathy Imbalance Hypothesis of Autism: A Theoretical Approach to Cognitive and Emotional Empathy in Autistic Development. The Psychological Record *2009 59:3*, 59(3), 489–510. 10.1007/BF03395675

Smith, S. D., McIver, T. A., Di Nella, M. S. J., & Crease, M. L. (2011a). The effects of valence and arousal on the emotional modulation of time perception: Evidence for multiple stages of processing. Emotion, 11(6), 1305–1313. 10.1037/A0026145

Smith, S. D., McIver, T. A., Di Nella, M. S. J., & Crease, M. L. (2011b). The effects of valence and arousal on the emotional modulation of time perception: evidence for multiple stages of processing. Emotion (Washington, D.C.), 11(6), 1305–1313. 10.1037/A0026145

Stupacher, J., Mikkelsen, J., & Vuust, P. (2022). Higher empathy is associated with stronger social bonding when moving together with music. Psychology of Music, 50(5), 1511–1526. 10.1177/03057356211050681

Suzuki, K., Lush, P., Seth, A. K., & Roseboom, W. (2019). Intentional Binding Without Intentional Action. Psychological Science, 30(6), 842–853. 10.1177/0956797619842191

Tamm, M., Uusberg, A., Allik, J., & Kreegipuu, K. (2014). Emotional modulation of attention affects time perception: Evidence from event-related potentials. Acta Psychologica, 149, 148–156. 10.1016/J.ACTPSY.2014.02.008

Trope, Y., & Liberman, N. (2010). Construal-Level Theory of Psychological Distance. Psychological Review. 10.1037/a0018963

Unruh, F., Landeck, M., Oberdörfer, S., Lugrin, J. L., & Latoschik, M. E. (2021). The Influence of Avatar Embodiment on Time Perception - Towards VR for Time-Based Therapy. Frontiers in Virtual Reality, 2, 658509. 10.3389/FRVIR.2021.658509/BIBTEX

Unruh, F., Vogel, D., Landeck, M., Lugrin, J. L., & Latoschik, M. E. (2023). Body and Time: Virtual Embodiment and its Effect on Time Perception. IEEE Transactions on Visualization and Computer Graphics, 29(5), 2626–2636. 10.1109/TVCG.2023.3247040

Vatakis, A., Allman, M. J., & Boston, L. |. (2015). Time Distortions in Mind. Time Distortions in Mind, 406. 10.26530/OAPEN_613387

Vicario, C. M., Scavone, V., Lucifora, C., Falzone, A., Pioggia, G., Gangemi, S., Craparo, G., & Martino, G. (2023). Evidence of abnormal scalar timing property in alexithymia. PLOS ONE, 18(1), e0278881. 10.1371/JOURNAL.PONE.0278881

Wallace, G. L., & Happé, F. (2008). Time perception in autism spectrum disorders. Research in Autism Spectrum Disorders, 2(3), 447–455. 10.1016/J.RASD.2007.09.005

Walsh, P. J. (2013). Empathy, Embodiment, and the Unity of Expression. Topoi *2013 33:1*, 33(1), 215–226. 10.1007/S11245-013-9201-Z

Walter, H. (2012). Social cognitive neuroscience of empathy: Concepts, circuits, and genes. Emotion Review, 4(1), 9–17. 10.1177/1754073911421379

Weiler, M., Acunzo, D. J., Cozzolino, P. J., & Greyson, B. (2024). Exploring the transformative potential of out-of-body experiences: A pathway to enhanced empathy. Neuroscience & Biobehavioral Reviews, 163, 105764. 10.1016/J.NEUBIOREV.2024.105764

Weng, C. C., Wang, N., Zhang, Y. H., Wang, J. Y., & Luo, F. (2022). The Effect of Electrical Stimulation–Induced Pain on Time Perception and Relationships to Pain-Related Emotional and Cognitive Factors: A Temporal Bisection Task and Questionnaire–Based Study. Frontiers in Psychology, 12, 800774. 10.3389/FPSYG.2021.800774/BIBTEX

Wilson, M. (2002). Six views of embodied cognition. Psychonomic Bulletin & Review *2002 9:4*, 9(4), 625–636. 10.3758/BF03196322

Wittmann, M. (2015). Modulations of the experience of self and time. Consciousness and Cognition, 38, 172–181. 10.1016/J.CONCOG.2015.06.008

Wohl, M. J. A., & McGrath, A. L. (2007). The Perception of Time Heals All Wounds: Temporal Distance Affects Willingness to Forgive Following an Interpersonal Transgression. Personality and Social Psychology Bulletin, 33(7), 1023–1035. 10.1177/0146167207301021

Yan, C., Wang, H., Jiang, X., & Wang, Z. (2025). Attention modulates subjective time perception across eye movements. Vision Research, 227, 108540. 10.1016/J.VISRES.2025.108540

Yang, Y., Zhao, J., Zhang, H., Bi, T., Tian, J., Li, Q., & Guo, C. (2024). The mutual influences between working memory and empathy for pain: the role of social distance. Social Cognitive and Affective Neuroscience, 19(1). 10.1093/SCAN/NSAE061

Zaki, J. (2014). Empathy: A motivated account. Psychological Bulletin.

Zaki, J. (2017). Moving beyond Stereotypes of Empathy. Trends in Cognitive Sciences, 21(2), 59–60. 10.1016/j.tics.2016.12.004

Zaki, J. (2020). Integrating empathy and interpersonal emotion regulation. Annual Review of Psychology, 71(1), 517–540. 10.1146/annurev-psych-010419-050830

Zaki, J., Bolger, N., & Ochsner, K. (2008). It takes two: The interpersonal nature of empathic accuracy: Research article. Psychological Science, 19(4), 399–404. 10.1111/J.1467-9280.2008.02099.X

